# Surf4 deficiency reduces intestinal lipid absorption and secretion and decreases metabolism in mice

**DOI:** 10.1101/2023.02.08.527677

**Authors:** Geru Tao, Hao Wang, Yishi Shen, Lei Zhai, Boyan Liu, Bingxiang Wang, Wei Chen, Sijie Xing, Yuan Chen, Hong-mei Gu, Shucun Qin, Da-wei Zhang

## Abstract

**Background:** Postprandial dyslipidemia is a causative risk factor for cardiovascular disease. The majority of absorbed dietary lipids are packaged into chylomicron and then delivered to circulation. Previous studies showed that Surfeit4 (Surf4) mediates very low-density lipoprotein secretion from hepatocytes. Silencing hepatic Surf4 markedly reduces the development of atherosclerosis in different mouse models of atherosclerosis without causing hepatic steatosis. However, the role of Surf4 in chylomicron secretion is unknown.

**Methods:** We developed inducible intestinal-specific *Surf4* knockdown mice (*Surf4*^IKO^) using *Vil*1Cre-ER^T2^ and *Surf4*^flox^ mice. Metabolic cages were used to monitor mouse metabolism. Enzymatic kits were employed to measure serum and tissue lipid levels. The expression of target genes was detected by qRT-PCR and Western Blot. Transmission electron microscopy and radiolabeled oleic acid were used to assess the structure of enterocytes and intestinal lipid absorption and secretion, respectively. Proteomics was performed to determine changes in protein expression in serum and jejunum.

**Results:** *Surf4*^IKO^ mice, especially male *Surf4*^IKO^ mice, displayed significant body weight loss, increased mortality, and reduced metabolism. *Surf4*^IKO^ mice exhibited lipid accumulation in enterocytes and impaired fat absorption and secretion. Lipid droplets and small lipid vacuoles were accumulated in the cytosol and the endoplasmic reticulum lumen of the enterocytes of *Surf4*^IKO^ mice, respectively. Surf4 colocalized with apoB and co-immunoprecipitated with apoB48 in differentiated Caco-2 cells. Intestinal Surf4 deficiency also significantly reduced serum triglyceride, cholesterol and free fatty acid levels in mice. Proteomics data revealed that diverse pathways were altered in *Surf4*^IKO^ mice. In addition, *Surf4*^IKO^ mice had mild liver damage, decreased liver size and weight, and reduced hepatic triglyceride levels.

**Conclusion:** Our findings demonstrate that intestinal Surf4 plays an essential role in lipid absorption and chylomicron secretion and suggest that the therapeutic use of Surf4 inhibition requires highly cell/tissue-specific targeting.

**Highlights:**

- Intestinal Surf4 deficiency reduces chylomicron secretion.
- Lack of intestinal Surf4 reduces absorption of dietary lipids.
- Silencing intestinal Surf4 reduces serum lipid levels.
- Intestinal Surf4 deficiency leads to lipid accumulation in intestinal villi.
- Intestinal Surf4 is essential for dietary lipid absorption and secretion.

## Introduction

Dietary lipids are an important determinant of circulating lipids that provide energy to various organs. However, increasing evidence suggests that elevated levels of postprandial lipids, including triglycerides (TG) and remnant cholesterol, induce endothelial dysfunction and increase risk for atherosclerotic cardiovascular disease, such as myocardial infarction and ischemic heart disease. People in Western societies are mostly in the fed state over the course of a day, which prolongs their exposure to high postprandial lipids, increasing the risk of cardiovascular events ^1, 2^. After a meal, lipid uptake occurs primarily in the intestine lumen, where lipids are emulsified by bile acids and hydrolyzed by various lipases ^1^. The majority of absorbed lipids are then esterified and packaged into lipoproteins in enterocytes, which are then released from the basolateral side of enterocytes, transported to the lymphatic system and then circulation through the thoracic duct ^3–5^.

Absorption and secretion of dietary lipids are regulated by diverse mechanisms ^5–8^. In enterocytes, the majority of dietary lipids are assembled into apolipoprotein B-48 (apoB-48) containing chylomicrons (CM). Microsomal triglyceride transfer protein (MTP) mediates and facilitates lipidation of apoB-48 protein co-translationally in the endoplasmic reticulum (ER) to generate dense apoB-48 containing lipoprotein particles, which are then fused with apoB-48-free lipid droplets in the ER lumen to form pre-CMs ^5, 7, 9–14^. Pre-CMs are too large to use the classic coat protein complex II (COPII) vesicles to exit the ER; instead, specific pre-CM transport vesicles (PCTV) transport pre-CMs from the ER to the Golgi, where pre-CMs are processed to mature CM and subsequently released from the basolateral side of enterocytes. PCTV-mediated ER export of pre-CMs is believed to be the rate-limiting step in transporting dietary lipids to circulation ^3–5, 7^. The formation of PCTV involves various proteins, such as liver fatty acid binding protein (FABP) ^15–17^, CD36 ^18, 19^, VAMP7 ^4, 20^, and v-SNARE ^21^. apoB-48, the main structural apolipoprotein on CM, is also required for PCTV budding from the ER ^22^. We and others reported that surfeit locus protein 4 (Surf4), a cargo receptor in the ER, interacts with apoB100 and mediates ER export of very low-density lipoprotein (VLDL) ^23, 24^. PCTV is larger than VLDL transport vesicles (VTV), and the mechanisms underlying the formation of PCTV and VTV are different ^5, 25, 26^. For example, Surf4 interacts with Sar1b, a GTPase and a component of COPII, to facilitate ER export of VLDL in hepatocytes. Sar1b also interacts with apoB48 ^4^. However, COPII components are required for ER export of VLDL in hepatocytes but not for the budding of PCTV from the ER in enterocytes; instead, COPII components mediate fusion of PCTV with the Golgi ^27, 28^. Whether a cargo receptor is required for ER export of pre-CM and the potential role of Surf4 in this process are unclear.

Surf4 is a cargo receptor located in ER membrane. It is ubiquitously expressed and can facilitate ER export of diverse secretory proteins ^29–32^. Surf4 directly interacts with apoB-100 and mediates VLDL secretion ^33, 34^. Hepatocyte-specific Surf4 knockout (*Surf4*^LKO^) mice have impaired VLDL secretion and significantly reduced plasma cholesterol, TG and apoB levels. Furthermore, knockdown of Surf4 expression significantly reduces the development of atherosclerosis in *Ldlr*^-/-^ and *apoE*^-/-^ mice ^35^. Here, to study its potential role in CM secretion, we used *Vil*1^Cre-ERT2^ mice to generate inducible intestinal-specific *Surf4* knockdown mice. We found that knockdown of Surf4 in the intestinal epithelium (*Surf4*^IKO^) significantly reduced intestinal fat absorption and secretion, as well as serum lipid levels, indicating an important role of intestinal Surf4 in dietary lipid metabolism.

## Material and methods

### Data availability

The authors declare that all supporting data are available within the article and its online supplementary files. The corresponding authors may provide additional data supporting the findings of this study upon reasonable request.

### Animals

*Surf4*^flox^ mice in C57BL/6J background were generated in Biocytogen (Beijing, China) ^23, 36^. Exon 2 of the *Surf4* gene was flanked by the loxP sites via CRISPR-Cas9 (Figure S1A). *Vil*1Cre-ER^T2^ mice were purchased from the Jackson Laboratory and bred with *Surf4*^flox^ mice to generate inducible intestine-specific *Surf4* knockdown mice. All mice used in the study were in the C57BL/6J background. Mice were housed in the facility at Shandong First Medical University (Taian, China) and fed *ad libitum* a standard laboratory rodent diet containing 20% protein, 5% fat, and 48.7% carbohydrates (Keao Xieli, Beijing, China, or LabDiet, PICO Laboratory Rodent Diet 20). Genotyping was conducted using PCR (Primers in Table S1). Cre recombinase expression was induced by intraperitoneal injection of tamoxifen (75 mg /kg/day in 100 μl corn oil for 4 consecutive days). All animal procedures were conducted following the guidelines of the Animal Care and Use Committee of Shandong First Medical University.

### Quantitative real-time PCR (qRT-PCR)

Total RNAs were extracted using TRIzol™ (Invitrogen). qPCR was conducted using TransStart Top Green qPCR SuperMix and specific primers (Table S1). Relative expression was calculated using 2^-ΔΔ^ method with *Gapdh* as the internal control.

### Western blot analysis

Tissue samples were lysed in RIPA buffer. Protein concentrations were determined with the BCA kit (Solarbio). Tissue homogenate or serum samples were applied to immunoblotting using specific antibodies (Table S2). Images were acquired on a chemiluminescence detection system (Clinx Science Instruments) and analyzed using the software Image J 1.53e. Densitometry was determined and used to quantify protein expression. Loading controls for immunoblotting of tissue homogenates (ACTIN or CALNEXIN) were quantified on the same membrane as the target proteins. Serum samples diluted 5-fold were run on a separated gel for immunoblotting of Albumin to obtain non-saturated exposures for quantification of serum proteins. Densitometry of targets was normalized to that of the loading control from the same mouse.

### Serum parameters

Blood samples were collected from mice (8-12 weeks old) via the tail vein and centrifuged at 3,000 rpm for 10 min to isolate serum for the measurement of serum levels of total cholesterol (TC), triglyceride (TG), high-density lipoprotein cholesterol (HDL-C), free fatty acid (FFA), ketone bodies, as well as aspartate aminotransferase (AST) and alanine aminotransferase (ALT) activity, using their specific colorimetric kits according to the manufacturer’s instructions. TC, TG and HDL-C kits were from BioSino Biotechnology and Science Inc, AST and ALT kits from Nanjing Jiancheng Bioengineering Institute, and ketone bodies from Solarbio (China). Fast protein liquid chromatography (FPLC) was performed on an AKTA system (GE, ÄKTA pure 25 L1).

### Histological analysis

Mouse jejunum was fixed in formalin and then subjected to H&E and Oil Red O staining (Wuhan Servicebio Technology Company, China). Sections were visualized under a microscope (Olympus, BX53). Intestinal tissue injury was evaluated from H&E stained sections following the criteria of Chiu’s score method ^37^.

### Transmission electron microscopy (TEM)

Male mice were fasted for 10 h and received an oral gavage of corn oil (10 μL/g). 2 h later, fresh jejunum samples were collected and fixed in glutaraldehyde, dehydrated, embedded and sectioned as described ^23^. Images were visualized in a Hitachi TEM system (HT7800).

### Tissue and feces lipid measurement

Lipids were extracted from tissues using the modified Folch method ^38, 39^. Tissue homogenate was mixed with chloroform: methanol (2:1). The lipid-containing phase was collected and mixed with chloroform containing 2% Trion X-100. Samples were dried under N_2_ and then redissolved in H_2_O. TG and TC were measured using their specific kits.

Fecal pellets were collected and weighed from each mouse over a 24-h period. Lipids were extracted using chloroform/methanol (2:1;v/v) as indicated ^40^. Lipid mass was weighed and then dissolved in ethanol. TC, TG and FFA were measured using their specific kits.

### Detection of intestinal triglyceride secretion

Mice were fasted for 10 h and received an intraperitoneal injection of tyloxapol (500 mg/kg). 30 min later, mice were given an oral gavage of corn oil (10 μL/g). Blood was collected at 0, 1, 2, 3, 4 h. Serum TG was measured using a commercial kit.

### Intestinal permeability assay

Mice were fasted for 12 h and received an oral gavage of PBS containing FITC-dextran (200 mg/kg) ^41^. 4 h later, blood samples were collected to measure fluorescence signal on a plate reader (Molecular Devices, Spectramax i3x; 485 nm excitation/530 nm emission).

### Uptake and metabolism of [^14^C] oleic acid

Male mice (10 weeks old, n=5) were fasted for 12 h and received an oral gavage of olive oil (10 μL/g) containing 0.1 μCi/g [^14^C]-oleic acid at MITRO Biotech Co. Ltd. (Nanjing, China). Blood samples were collected at 30, 60, 120, 180 and 240 min after gavage. Tissues were collected at the endpoint. [^14^C] levels were measured using scintillation counting (ALOKA, LSC-8000).

### Metabolic profiling

Male and female mice (9-10 weeks old, n=5) were housed individually and monitored undisturbed for 3 days by a Comprehensive Lab Animal Monitoring System (Oxymax®-CLAMS, Columbus Instruments). Data from the last 48 h were used for analysis.

### Confocal and Co-immunoprecipitation

Caco-2 cells were maintained in DMEM containing 10% (v/v) FBS and differentiated as described in ^42^. Cells were plated on a 100 mm dish or coverslip and maintained for 21 days after confluency to induce differentiation. Cells were then incubated in DMEM containing 0.4 mM oleic acid for 4 h, followed by immunoprecipitation or confocal microscopy.

Immunoprecipitation was performed as described ^43, 44^. Briefly, cells were lysed in lysis buffer A (1% Triton, 150 mM NaCl, 50 mM HEPES, pH 7.4) containing 1× Complete EDTA-free protease inhibitors. Equal amount of total protein was applied to a rabbit anti-Surf4 or purified rabbit IgG and protein G beads. Immunoprecipitated proteins were eluted from the beads using 2 × SDS-PAGE sample buffer and then applied to immunoblotting.

Confocal microscopy was conducted as described ^45–47^. Cells were fixed and permeabilized with cold methanol and then incubated with a goat anti-apoB polyclonal antibody and a rabbit anti-Surf4 antibody. Antibody binding was detected using Alexa Fluor 568 donkey anti-rabbit IgG and Alexa Fluor 488 donkey anti-goat IgG. Localizations of apoB and Surf4 were determined using a Leica SP5 laser scanning confocal microscope.

### Proteomic and bioinformatic analysis

Serum and jejunum (male, 9-10 weeks old, n=5) were collected from euthanized mice and shipped to Jingjie PTM-Biolab for proteomic analysis. Samples were subjected to standard protein extraction, trypsin digestion and liquid chromatography-mass spectrometry (LC-MS). False discovery rate was adjusted to < 1%. Differentially expressed proteins were identified when the FoldChange was more than 1.5 and adjusted *p* value was less than 0.05. Kyoto Encyclopedia of Genes and Genomes (KEGG) database was used to annotate protein pathways. The online tool of Jingjie PTM-BIO Shiny Tool was used to analyze data and generate graphics.

### Statistical analysis

All statistical analyses were performed using Prism 9. The Mann-Whitney test was used to determine significant differences between the two groups, and the Kruskal-Wallis test followed by a Dunn’s test was used to determine the significance among different groups. All data were presented as median with 95% confidence limits. The significance was defined as *p*<0.05.

## Results

### Generation of intestine-specific Surf4 knockdown mice

To generate an inducible intestine-specific Surf4 knockdown mice, we bred *Surf4*^flox^ mice with mice carrying tamoxifen-inducible Cre recombinase under the control of an intestinal epithelium-specific villin 1 (*Vil1*) promotor (*Vil*1Cre-ER^T2^) (Figure S1A). Heterozygous *Vil*1Cre-*Surf4*^flox^ mice were bred to generate homozygous mice, which were confirmed by the presence of loxP sites and Cre (Figure S1B). The resulting mice were fertile and indistinguishable from *Surf4*^flox^ mice. Exon 2 of the *Surf4* gene in *Surf4*^flox^ mice was flanked with LoxP sites. After tamoxifen administration, Cre-mediated recombination of the Surf4 gene deleted amino acids Phe^17^ to Leu^78^ and introduced a stop codon after Phe^17^ to result in homozygous (*Surf4*^IKO^) and heterozygous (*Surf4*^HTZ^) intestinal epithelium-specific Surf4 knockdown mice. As shown in Figures 1A and B, Surf4 expression was markedly reduced in the jejunal scrapings of both male and female *Surf4*^IKO^ mice but not *Surf4*^HTZ^ mice, compared to *Surf4*^flox^ mice. qRT-PCR data confirmed that Surf4 expression in *Surf4*^IKO^ mice was reduced in the intestine. We observed higher residual mRNA levels of Surf4 in female *Surf4*^IKO^ mice than in male *Surf4*^IKO^ mice (Figures S1C and S1D). Female *Surf4*^IKO^ mice also seemed to have a higher residual level of Surf4 protein than male *Surf4*^IKO^ mice (Figures 1A and 1B). A detailed analysis, however, showed that the difference did not reach statistical difference (mean expression of Surf4 protein =0.133 in male and 0.233 in female *Surf4*^IKO^, p=0.092). Furthermore, the reduction in Surf4 protein levels was greatly significant in both male and female *Surf4*^IKO^ mice compared to the corresponding *Surf4*^flox^ mice (mean expression of Surf4 protein =0.9830 in male and 0.7956 in female *Surf4*^flox^ mice, respectively). qRT-PCR data showed that mRNA levels of Surf4 were not changed in the heart, liver, spleen, lung, kidney, and adrenal gland in both male and female mice, or the ovary in female mice. However, male *Surf4*^IKO^ mice showed a mild but significant reduction in the mRNA levels of Surf4 in the testis (Figures S1C and S1D). Therefore, Surf4 expression is selectively and markedly reduced in the intestine of *Surf4*^IKO^ mice.

**Figure 1.**
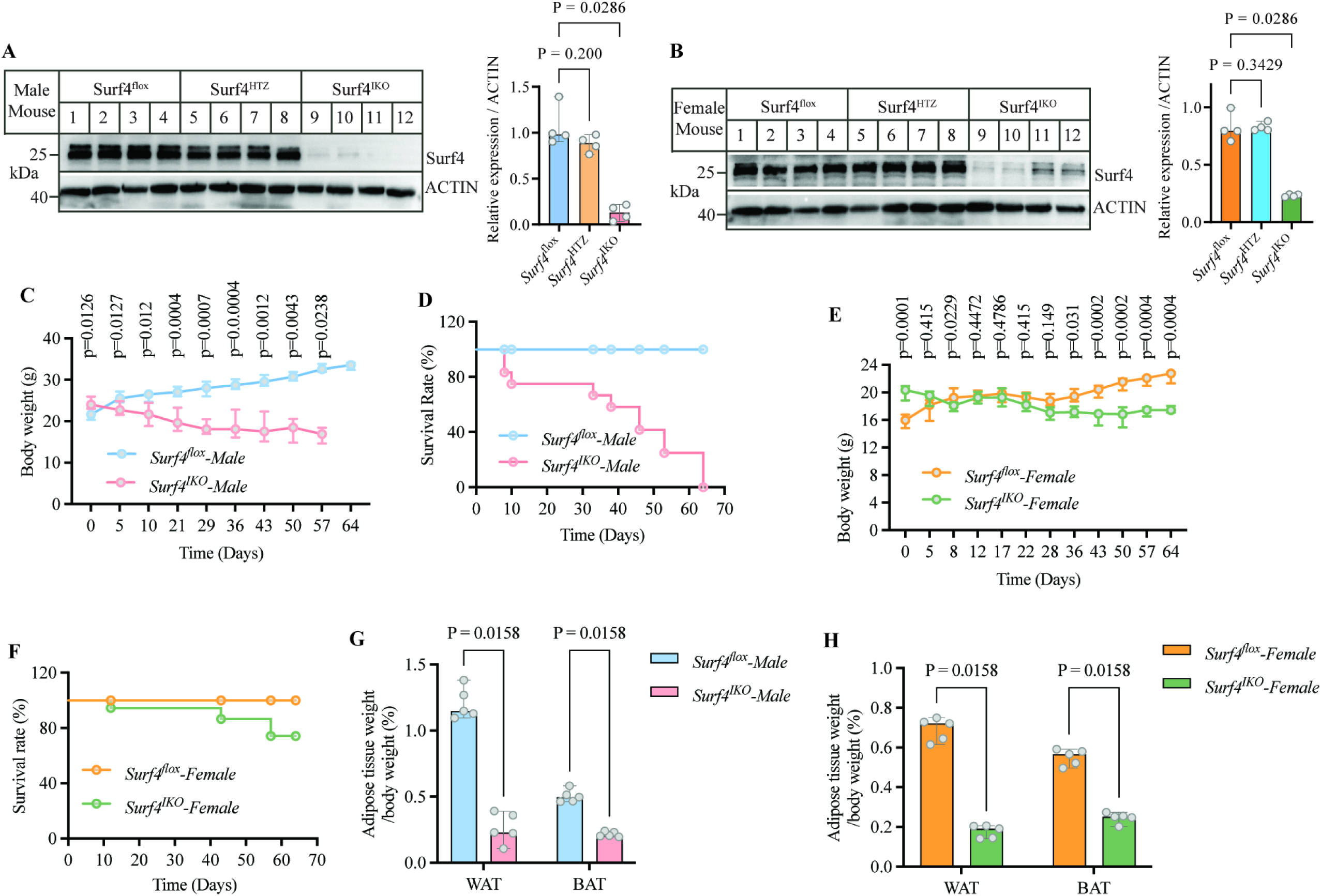
Generation of *Surf4*^IKO^ mice. **A and B**. Western blot. Male (A) and female (B) *Surf4*^flox^, *Surf4*^HTZ^ and *Surf4*^IKO^ mice (8-12-week-old, n=4, biological replicates) were injected with tamoxifen for 4 days and then fed a standard laboratory rodent diet for 4 days. Jejunum scrapings were collected and subjected to immunoblotting. ACTIN was the loading control. **C to F.** Phenotype of *Surf4*^IKO^ mice (8-12-week-old, 6 males and 6 females of *Surf4*^flox^ mice and 12 males and 12 females of *Surf4*^IKO^ mice, biological replicates). *Surf4*^flox^ and *Surf4*^IKO^ mice were injected with tamoxifen for 4 days and then fed a standard laboratory rodent diet. Body weight and survival rate of male (C and D) and female (E and F) *Surf4*^flox^ and *Surf4*^IKO^ mice starting at the beginning of injection (Day 0). The survival experiments in panels D and F were performed once. **G and H**. Adipose tissue weight (n=5, biological replicates). Epididymal white adipose tissue (eWAT) and interscapular brown adipose tissue (BAT) were dissected from male (G) and female (H) mice and immediately weighed. The percentages of eWAT and BAT were relative to the body weight of the same mouse. The Mann-Whitney test was used to determine significant differences between two groups.

*Vil*1Cre-*Surf4*^flox^ mice injected with corn oil exhibited a comparable body weight as *Surf4*^flox^ mice (Figure S2), while male *Surf4*^IKO^ mice lost significant body weight and appeared to have approximately 70% survival at day 30 after tamoxifen administration. Their health then deteriorated rapidly, and all mice reached a humane endpoint at 64 days (Figures 1C and D). On the other hand, the body weight of female *Surf4*^IKO^ mice remained stable 28 days after tamoxifen injection and then began to decrease, but to a much lower degree than that of male mice (Figure 1E). They also had a higher survival rate of about 80% at 64 days (Figure 1F). Detailed analysis showed that both male and female *Surf4*^IKO^ mice had a significant reduction in both white (WAT) and brown adipose tissue (BAT) weights only 4 days after completion of tamoxifen administration (Figures 1G and H). Thus, intestinal-specific knockdown of Surf4 results in weight loss and increased mortality in mice, especially in male mice.

### Effects of intestinal Surf4 deficiency on mouse metabolism

We next determined the effect of intestinal *Surf4* knockdown on mouse metabolism using metabolic cages. Male *Surf4*^IKO^ mice consumed more food in the light cycle but consumed significantly less food and water in the dark and 24 h cycles than male *Surf4*^flox^ mice (Figures 2A and B). In *Surf4*^IKO^ mice, heat production was significantly reduced in the light cycles and trended downward in the dark and 24 h cycles (Figure 2C), and activity was reduced in the dark and 24 h cycles (Figure 2D). We also observed that *Surf4*^IKO^ mice had significantly reduced oxygen consumption in all cycles (VO_2_; Figures 2E and S3A) and carbon dioxide release in the light and 24 h cycles (VCO_2_; Figures 2F and S3B). Respiratory quotient (RQ) was significantly increased in male *Surf4*^IKO^ mice in the dark and 24 h cycles (Figures 2G and S3C), indicating that they used less fat in these cycles. As shown in Figures 2H-N and S3D-F, female *Surf4*^IKO^ mice exhibited similar trends, decreased food and water intake, heat production, activity, and VO_2_, and increased RQ. Changes in food intake in the dark cycle (Figure 2H), water consumption in the light and 24-h cycles (Figure 2I), and RQ in the light and 24-h cycles (Figure 2N) were statistically significant, while other changes did not reach statistical significance. Therefore, intestinal-specific knockdown of Surf4 significantly reduces metabolism in mice, with a more pronounced effect in male mice.

**Figure 2.**
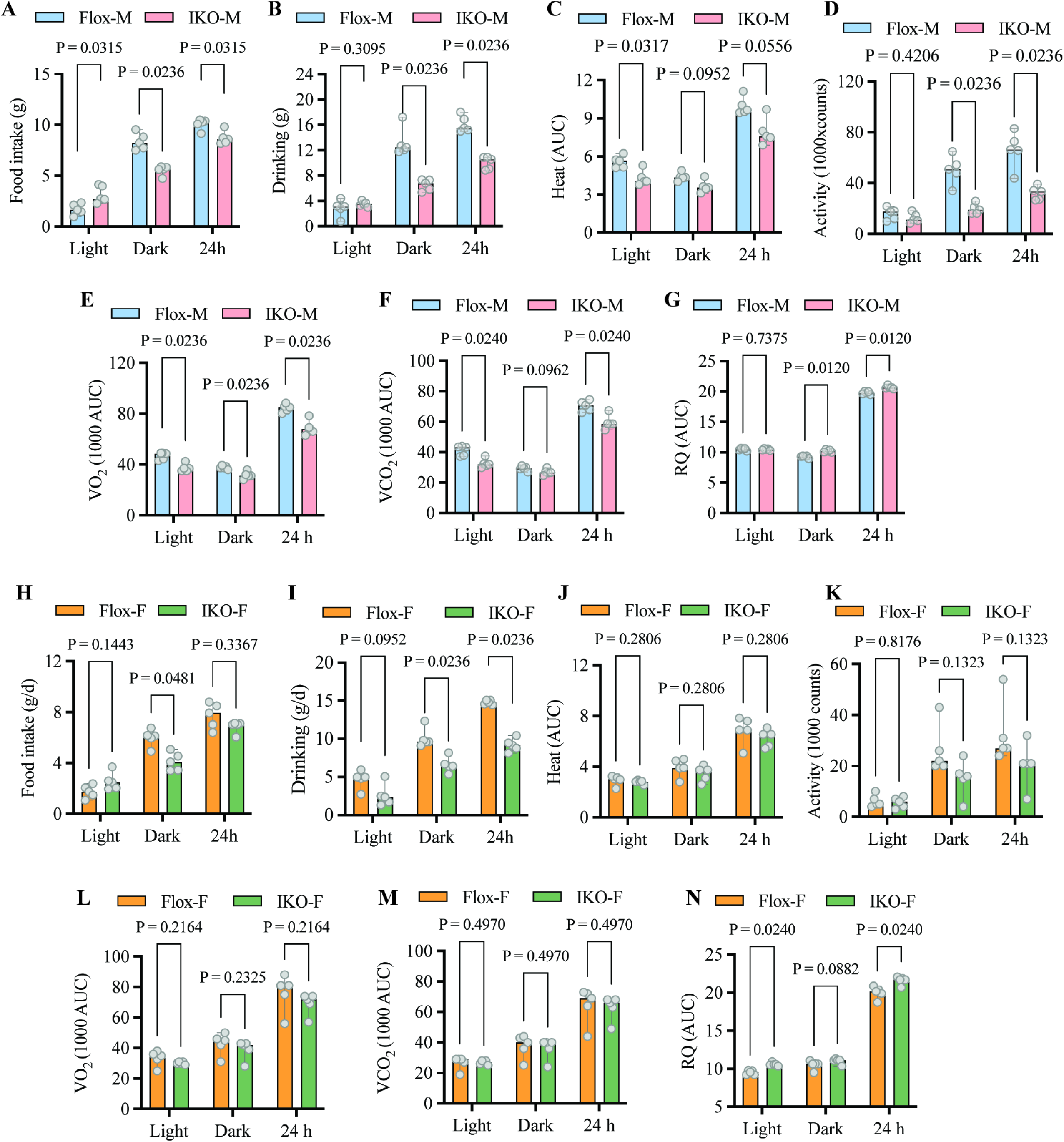
Effect of intestinal *Surf4* knockdown on mouse metabolism. Male and female *Surf4*^flox^ and *Surf4*^IKO^ mice (9 weeks old, n=5, biological replicates) were injected with tamoxifen for 4 days, fed a standard laboratory rodent diet for 4 days, and then subjected to metabolic cages for 72 h. **A-G.** Male mice. **H-N**. Female mice. Food intake (A and H), drinking water (B and I), heat production (C and J), activity (D and K), VO_2_ (E and L), VCO_2_ (F and M), and RQ (G and N). Area under curve (AUC) was calculated using Prizm 9. The Mann-Whitney test was used to determine significant differences between two groups.

### Lipid accumulation in *Surf4* ^IKO^ mice intestine

We then analyzed the effect of Surf4 deficiency on the intestine. *Surf4*^IKO^ but not *Surf4*^HTZ^ mice displayed increased intestinal length compared to *Surf4*^flox^ mice (Figures 3A to C). The entire small intestine of *Surf4*^IKO^ mice looked whiter than that of *Surf4*^HTZ^ and *Surf4*^flox^ mice even after intestinal contents were flushed out (Figures 3A and B), suggesting intestinal lipid accumulation. Indeed, H&E and Oil red O staining showed accumulation of lipid droplets in villous enterocytes of both male (Figure 3D) and female *Surf4*^IKO^ mice (Figure S4A). To further confirm intestinal lipid accumulation, we measured intestinal TG and TC levels and found that they were significantly increased in the intestine of male and female *Surf4*^IKO^ mice (Figures 3E, 3F, S4B and S4C). On the other hand, *Surf4*^HTZ^ mice exhibited comparable intestinal TG and TC levels as *Surf4*^Flox^ mice. We then assessed post-fat feeding serum TG levels in the presence of tyloxapol, which inhibits lipoprotein lipase activity ^48^. As shown in Figures 3G and S4D, serum TG levels were increased dramatically 2 h after oral administration of corn oil and continued to increase slowly over 4 h in both male and female *Surf4*^flox^ mice. On the other hand, post-fat feeding serum TG levels in male and female *Surf4*^IKO^ mice were barely increased over the 4-h period. Consistently, analysis of area under curve (AUC) revealed a significant reduction in serum TG levels in Surf4 deficient mice. In addition, we observed that intestinal epithelial cells of *Surf4*^flox^ mice were tightly connected to each other by the desmosomal junctional apparatus near the lumenal brush border side (grade 0) and displayed only a few *Gruenhagen’s* space. In contrast, the jejunum of *Surf4* ^IKO^ mice showed extensive subepithelial spaces (Figures 3D and H). Consistently, the expression of the tight junction-associated protein, occludin and E-cadherin, was reduced in the jejunum of *Surf4*^IKO^ mice (Figure 3I), indicating intestine injury. We then used 4-kDa FITC-dextran to determine whether Surf4 deficiency severely disrupted intestinal integrity. Mice were fasted for 12 h prior to administration of FITC-dextran to minimize interference of intestinal contents on FITC-dextran absorption. Serum FITC-dextran levels were significantly lower in *Surf4*^IKO^ mice 4 h after oral gavage (Figure 3J), indicating reduced paracellular permeability of the intestinal epithelium or decreased absorption of FITC-dextran. Nevertheless, this finding suggests that intestinal Surf4 deficiency causes intestinal damage but does not result in notably leaky intestines, and the lack of intestinal Surf4 appears to impair CM release from enterocytes, leading to lipid accumulation.

**Figure 3.**
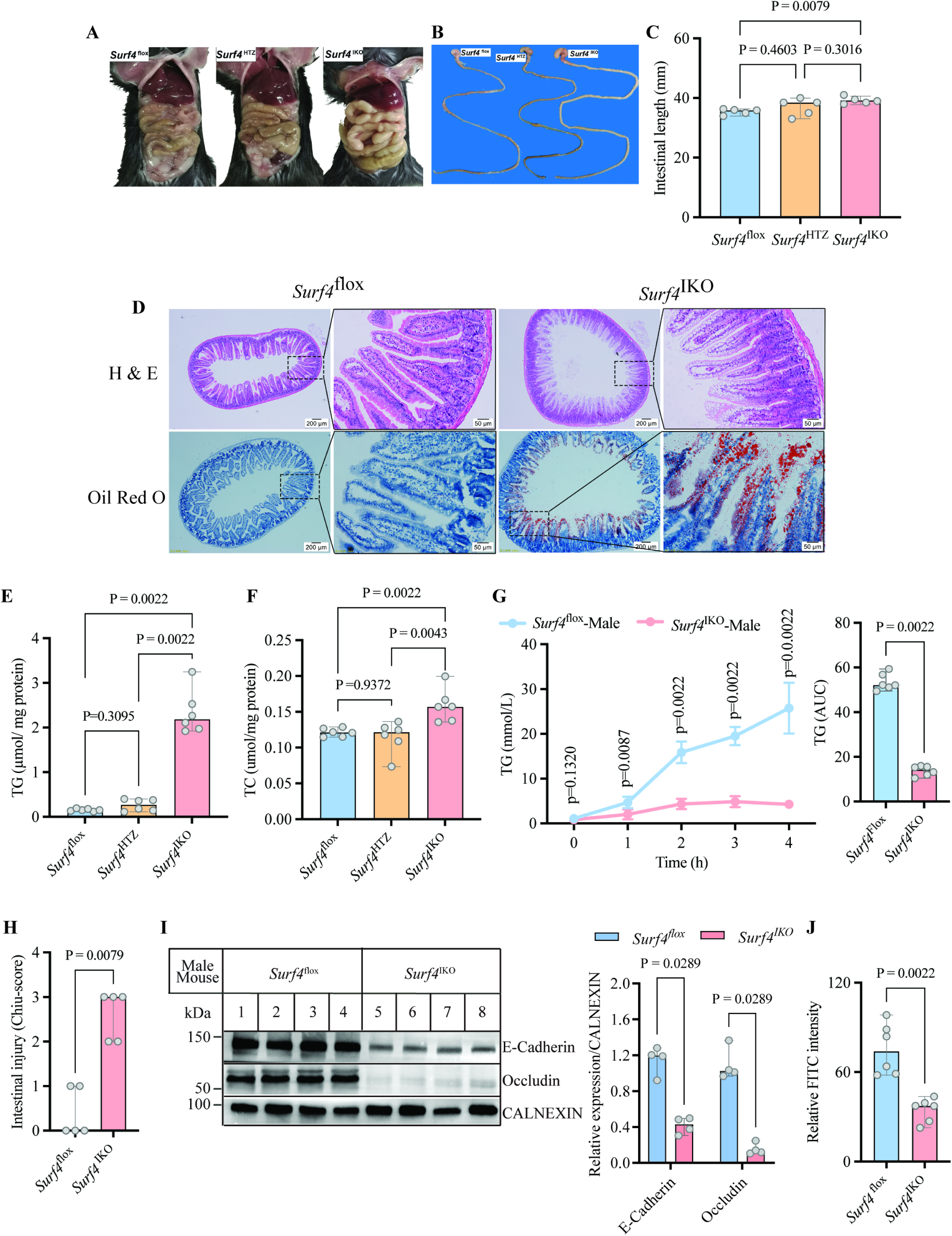
Lipid accumulation in *Surf4*^IKO^ mice intestines. Male *Surf4*^flox^, *Surf4*^HTZ^ and *Surf4*^IKO^ mice (9-10 weeks old) were injected with tamoxifen for 4 days and fed a standard laboratory rodent diet for 4 days before being subjected to euthanasia. **A.** Small intestines of *Surf4*^flox^, *Surf4*^HTZ^ and *Surf4*^IKO^ mice. Representative images were shown. Similar phenotypes were observed in other mice. **B and C.** Length of small intestines (n=5, biological replicates). **D.** H&E and Oil Red O staining of jejunum of *Surf4*^flox^ and *Surf4*^IKO^ mice (n=5, biological replicates). Representative images were shown. Similar results were observed in other mice. **E and F**. Jejunum lipids (n=6, biological replicates). Lipids were extracted from jejunum. TG (E) and TC (F) were measured and normalized to protein concentrations. **G.** Serum TG (n=6, biological replicates). Mice were fasted and received tyloxapol (IP) and an oral gavage of corn oil. Blood samples were collected at the indicated time. Serum TG was measured. AUC of serum TG levels of each mouse was calculated using Prizm 9. **H.** Intestinal tissue injury evaluated from H&E sections according to the criteria of Chiu’s score. **I.** Immunoblotting. Jejunum was collected from male *Surf4*^flox^ and *Surf4*^IKO^ mice (n=4, biological replicates). Equal amount of homogenate was subjected to immunoblotting using antibodies as indicated. Relative expression was the ratio of the densitometry of the target to that of CALNEXIN from the same mouse. **J.** Intestinal permeability using FITC-dextran (4 kDa). Serum fluorescence in pre- and 4 h post-gavage was measured and calculated. The Mann-Whitney test was used to determine significant differences between two groups.

### Reduced intestinal absorption and release of lipids in *Surf4* ^IKO^ mice

We then performed TEM on jejunum of mice fasted for 10 h and then orally gavaged with corn oil. We observed CM-like vacuoles (diameter: 249±27.893 nm) in the villi of *Surf4*^flox^ mice (Figure 4A). Conversely, Surf4-deficient villi displayed many small pre-CM (p-CM)-like vacuoles (diameter: 103.57±17 nm) in the ER lumen (RER) (Figure 4B), suggesting an important role for Surf4 in CM secretion. Given the critical role of apoB48 in CM assembly and secretion, we investigated the interaction between Surf4 and apoB48. We first conducted confocal microscopy in differentiated human enterocyte-like Caco-2 cells treated with oleate. Surf4 and apoB were detected by their specific antibody and shown in red and green fluorescence, respectively (Figure 4C). We observed yellow puncta and partial overlap of the two colour (Figures 4C and E), indicating partial colocalization of Surf4 and apoB. We also checked Surf4 and apoB in Caco-2 cells in the absence of oleic acid treatment and found that colocalization of Surf4 and apoB was significantly reduced compared to oleic acid-treated cells (Figures 4D and E). Oleic acid stimulates apoB secretion, which may enhance interaction and colocalization of Surf4 and apoB in the secretory pathway. We then performed co-immunoprecipitation experiments. A rabbit anti-Surf4 polyclonal antibody but not control rabbit IgG pulled down Surf4 from whole cell lysate (Figure 4F, lane 3 vs. 2). Caco-2 cells express apoB-100 and apoB-48, both of which co-immunoprecipitated with Surf4 (Figure 4F, lane 3). Taken together, these findings suggest that Surf4 is associated with apoB48 and mediates CM secretion.

**Figure 4.**
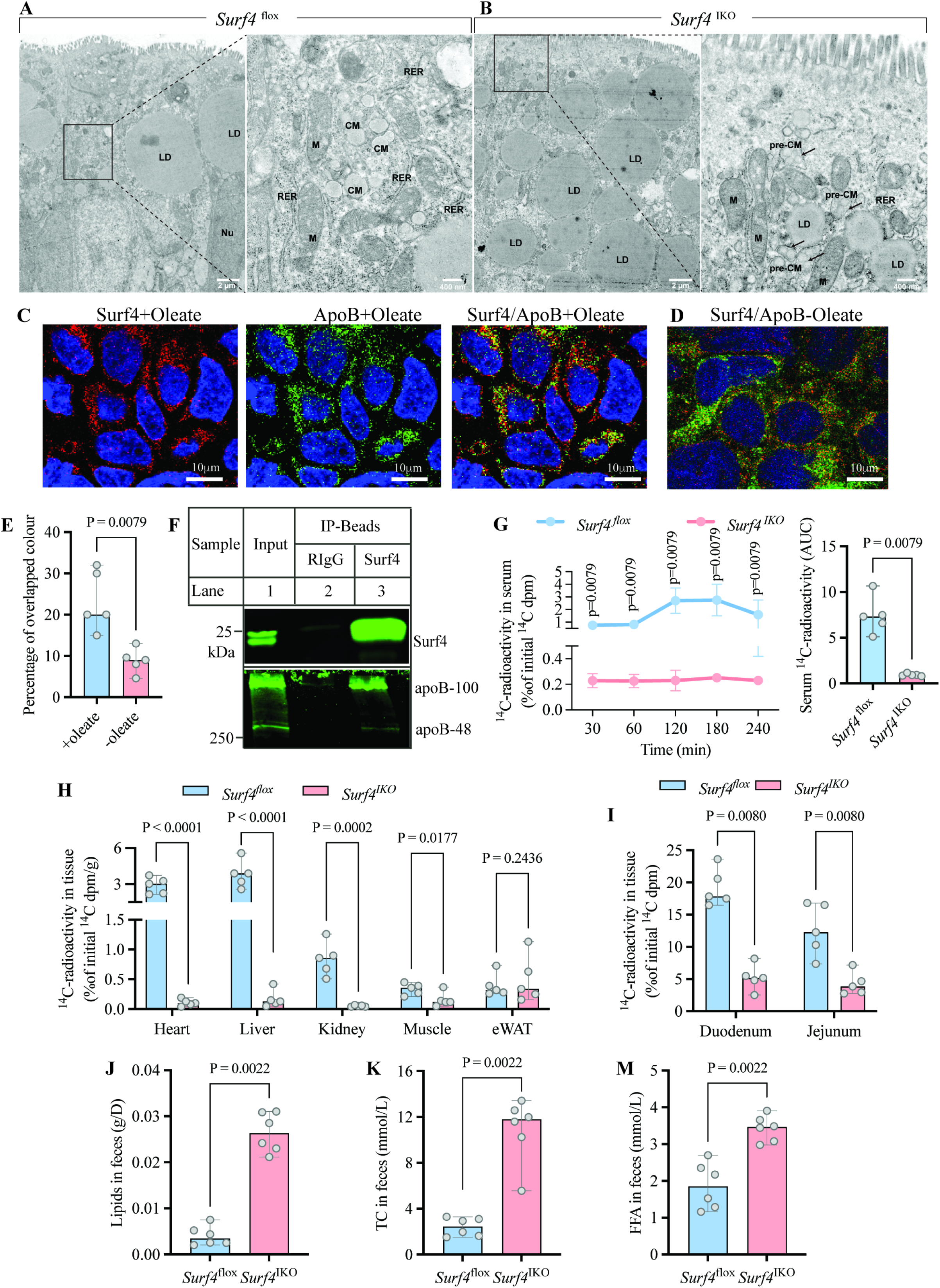
Reduced intestinal lipid absorption and release in *Surf4*^IKO^ mice. *Surf4*^flox^ and *Surf4*^IKO^ mice (9-week-old, male) were injected with tamoxifen for 4 days, fed a standard laboratory rodent diet for 4 days, and then treated as described below. **A and B**. TEM. Mice (n=3, biological replicates) were fasted for 10 hours and then received a bolus of 10 μL/g corn oil. 2 h later, jejunum was collected and subjected to TEM. Representative images were shown. Similar phenotypes were observed in other samples. **C to E.** Confocal microscopy. Caco2 cells treated with (C) or without (D) oleate conjugated to BSA (0.4 mM) for 4 h, fixed and permeabilized. Surf4 and apoB were detected by a rabbit anti-Surf4 (red) and a goat anti-apoB (green) antibody. Nuclei were visualized with DAPI and shown in blue (magnification: 100X). Color was quantified in ImageJ, and percentage of overlapped color (yellow) was calculated (E, n=5, technical replicates). **F.** Immunoprecipitation. Equal amount of whole cell lysate from Caco2 cells was incubated with purified rabbit IgG (RIgG) and a rabbit anti-Surf4 antibody. Whole cell lysate (Input) and immunoprecipitated proteins (IP-beads) were subjected to immunoblotting. Similar results were observed in three independent experiments (biological replicates). **G-I.** [^14^C] oleic acid absorption. Male mice (n=5, biological replicates) were fasted for 10 hours, followed by an oral gavage of 10 μL/g olive oil containing [^14^C] oleic acid. Blood samples were collected at the indicated times to measure serum [^14^C] activity (G). AUC of serum [^14^C] activity of each mouse was calculated using Prizm 9. Mice were euthanized at the endpoint (240 min). [^14^C] appearance in the heart, liver, kidney, muscle and eWAT (H), as well as in duodenum and jejunum (I) was measured. **J-K.** Fecal lipids. Feces were collected from male mice housed individually during 24 h (n=6, biological replicates). Lipids were extracted and weighed (J), TC (K) and FFA (M) were extracted and measured. The Mann-Whitney test was used to determine significant differences between two groups.

Next, we used [^14^C]-oleic acid to test the impact of Surf4 silencing on fat absorption and release. Mice were fasted for 10 h and then received a bolus of corn oil containing [^14^C]-oleic acid. Serum ^14^C radioactivity in *Surf4*^flox^ mice began to increase at 1 h, continued to increase markedly up to 2 h, remained stable at 3 h, and then began to decline (Figure 4G). On the other hand, *Surf4*^IKO^ mice exhibited flat and significantly low serum ^14^C-activity over the 4 h period. Consistently, AUC analysis revealed significantly reduced serum ^14^C-activity in intestinal Surf4-deficient mice. *Surf4*^IKO^ mice also had significantly lower levels of ^14^C-activity in the heart, liver, kidney and muscle than *Surf4*^flox^ mice, but ^14^C-activity in eWAT was not statistically different between the two genotypes (Figure 4H). Data were expressed as a percentage of ^14^C activity in each tissue to the initial total ^14^C activity, which was then normalized to the corresponding tissue weight. ^14^C activity in eWAT of *Surf4*^IKO^ and *Surf4*^flox^ mice was 397.2 and 1130.8, respectively. *Surf4*^IKO^ mice had significantly low eWAT weight compared to *Surf4*^flox^ mice (0.0612 g vs 0.1174 g, p=0.0119), consistent with the findings in Figures 1G and H. The significant reduction in eWAT weight of *Surf4*^IKO^ mice might affect ^14^C-oleic acid absorption. More studies are needed to understand the underlying cause. In addition, ^14^C radioactivity in the duodenum and jejunum of *Surf4*^IKO^ mice was significantly lower than that in *Surf4*^flox^ mice (Fig. 4I). These findings suggest that lipid absorption and secretion are impaired in *Surf4*^IKO^ mice. To confirm this possibility, we measured fecal lipids and found that total lipids, TC, and free fatty acids (FFA) levels were all significantly increased in both male (Figures 4J to M) and female *Surf4*^IKO^ mice (Figures S4E to G). In conclusion, *Surf4*^IKO^ mice have impaired dietary lipid absorption and secretion.

### Effect of Surf4 deficiency on intestinal protein expression

To explore how lipid metabolism was affected in jejunum of *Surf4*^IKO^ mice, we collected jejunum for Western blotting of proteins involved in cholesterol and fatty acid metabolism. As shown in Figures 5A and B, the protein levels of apoB-48, CD36, and MTP were comparable in the two genotypes, whereas apoA-I and apoE levels were significantly increased in *Surf4*^IKO^ mice. Conversely, we observed a significant reduction in the protein levels of apoA-IV, FABP2 (fatty acid transporters), NPC1L1 (cholesterol absorption), HMGCR (the rate-limiting enzyme of cholesterol biosynthesis), and MOGAT2 (diacylglycerol biosynthesis) in *Surf4*^IKO^ mice. We then determined the mRNA levels of these genes. Consistent with their reduced protein levels, the mRNA levels of *Apoa-4*, *Fabp2*, and *Npc1l1* were significantly reduced in *Surf4*^IKO^ mice (Figure 5C). Conversely, the mRNA levels of *Mogat2* and *Mttp* were comparable in *Surf4*^flox^ and *Surf4*^IKO^ mice, even though MOGAT2 protein levels were significantly reduced in *Surf4*^IKO^ mice. On the other hand, the mRNA levels of *Cd36* were significantly reduced, but its protein levels were unchanged in *Surf4*^IKO^ mice. The mRNA and protein levels of *Hmgcr* were significantly increased and decreased, respectively, in *Surf4*^IKO^ mice. Interestingly, we found that the mRNA levels of *Apob*, *Apoa1* and *Apoe* were all significantly reduced, although their protein levels were unchanged or even increased in *Surf4*^IKO^ mice. It was possible that the expression of these apolipoproteins was reduced at the transcriptional level, but their translation may be affected, resulting in no change or even an increase in their protein levels. Overall, these data suggest that *Surf4*^IKO^ mice may have reduced lipid absorption and synthesis in the jejunum to alleviate lipid accumulation.

**Figure 5.**
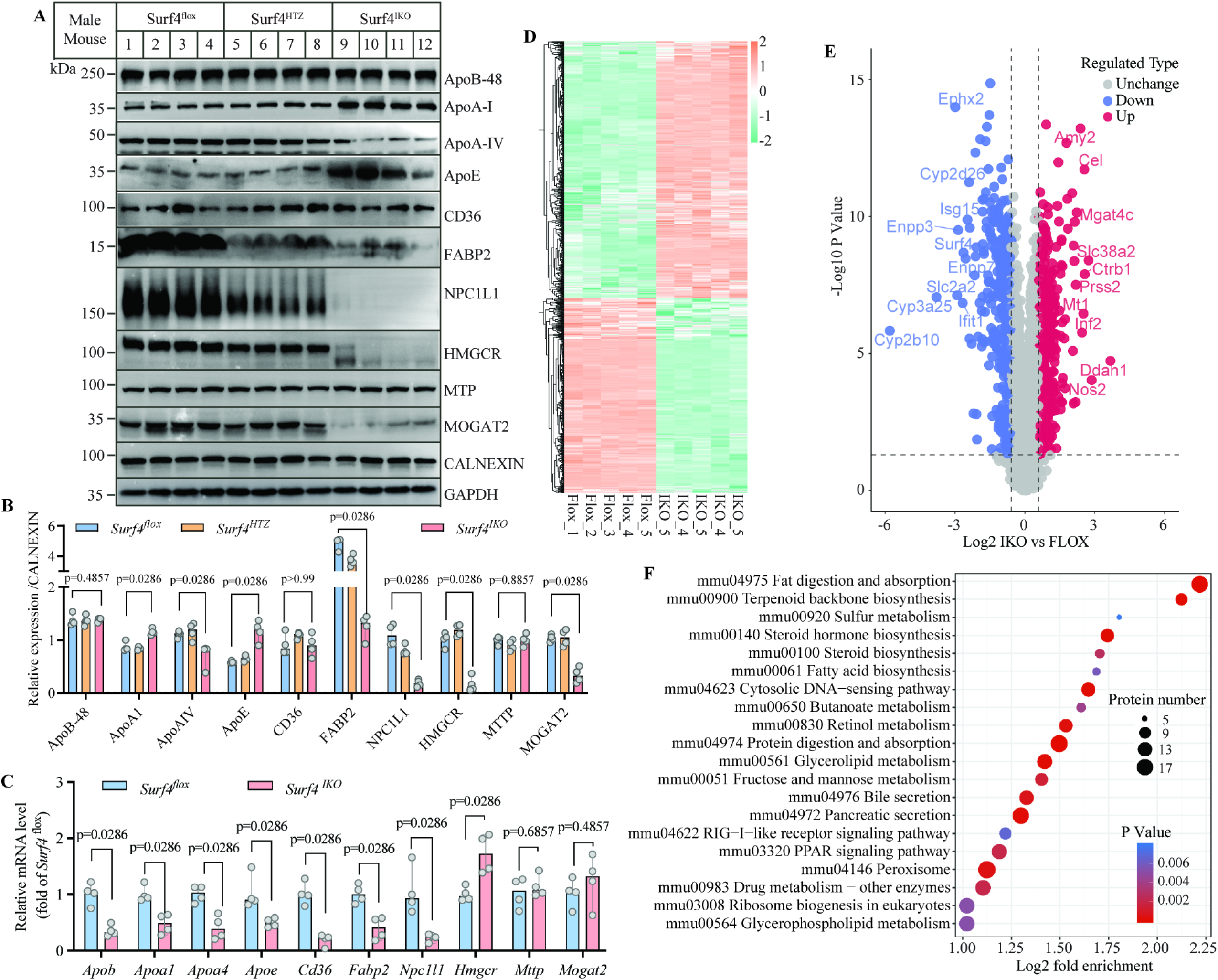
Effects of intestinal Surf4 deficiency on gene expression. **A and B.** Immunoblotting. Jejunum was collected from *Surf4*^flox^, *Surf4*^HTZ^ and *Surf4*^IKO^ mice. Equal amount of homogenate was subjected to immunoblotting using antibodies as indicated (A). Relative expression was the ratio of the densitometry of the target to that of CALNEXIN (B), which levels were comparable to that of GAPDH (A). **C.** qRT-PCR. Total RNA was extracted from jejunum of *Surf4*^flox^ and *Surf4*^IKO^ mice (n=4, biological replicates) for qRT-PCR. Relative expression was calculated using 2^-ΔΔ^ method with *Gapdh* as the internal control. **D-F.** Proteomics of mouse jejunum (n=5, biological replicates). *Surf4*^flox^ and *Surf4*^IKO^ mice (9-week-old, male) were injected with tamoxifen for 4 days, fed a standard laboratory rodent diet for 4 days. Jejunum was collected for proteomics. MaxQuant search engine (v.1.6.15.0) was used to identify jejunum proteins Heat map (D), volcano plot (E), and KEGG pathway enrichment (F). In heatmaps, each block represents a certain protein, and the horizontal axis represents each sample. Heatmap presents differentially expressed proteins. Values were z-score normalized, log2 label-free quantification intensity. The color gradient is from green (low values) to orange (high values). Proteomic data were analyzed, and graphic were generated by online system of PTM-Bio Shiny Tool. The Mann-Whitney test was used to determine significant differences between two groups.

In addition to absorption and secretion, TG pool in enterocytes is regulated by *de novo* lipogenesis and fatty acid oxidation. Therefore, we assessed expression of proteins important for these biological processes. As shown in Figure S5, the protein and mRNA levels of genes critical for lipogenesis (*Scd1*, *Fasn*, and *Acaca*) and fatty acid oxidation (*Cpt1a*) were significantly increased in the intestine of *Surf4*^IKO^, suggesting increased *de novo* lipogenesis and fatty acid oxidation.

We further performed proteomics on jejunum samples from male mice. A total of 61191 peptides were identified, matching 58916 unique proteins. Of these, 5373 quantifiable proteins were identified, including 497 upregulated and 370 downregulated proteins (Figures 5D and 5E, Tables S3 and S4). Bioinformatic enrichment analysis revealed that the most affected and enriched pathway was fat digestion and absorption (Figure 5F). Proteins with altered expression levels were also enriched in other pathways involved in digestion, such as protein digestion and absorption and pancreatic secretion. The 497 upregulated proteins were involved in a variety of biological processes (Figure S6A, Table S3), of which 97 proteins were associated with metabolism, including 29 in lipid transport and metabolism. 52 were associated with posttranslational modifications, protein turnover and chaperons, suggesting that lipid accumulation in the intestinal ER of *Surf4*^IKO^ mice may cause ER stress. The top two upregulated proteins were dimethylarginine dimethyaminohydrolase-1 (DDAH1, IKO/Flox=12, p=1.9E-05) and nitric oxide synthase 2 (NOS2, IKO/Flox=7, p=9.4E-05) (Table S3). DDAH1 and NOS2 can increase bioavailable NO to protect the gastrointestinal tract from injury and damage ^49, 50^. We also observed a dramatic increase in SLC38A2 (6.6-fold, p=4.0E-09), inverted formin 2 (INF2, 5.4-fold, p=1.7E-06), metallothionein 1A (MT1, 5.7-fold, p=3.5E-07), chymotrypsinogen B1 (CTRB1, 5.9-fold, p=1.3E-08), carboxyl ester lipase (CEL, 5.8-fold, p=1.9E-12), and amylase alpha 2A (AMY2, 5.2-fold, p=6.1E-14). SLC38A2 is an amino acid transporter that prefers short-chain neutral amino acid residues ^51^. INF2 regulates polymerization and depolymerization of actin filaments ^52^. MT1 acts as an anti-oxidant to detoxify heavy metals and prevent stress-induced cell damage ^53^. CTRB1, CEL and AMY2 are pancreatic serine proteinase, lipase, and amylase, respectively. These findings suggest that *Surf4*^IKO^ mice may develop mechanisms to mitigate intestinal damage.

The 370 downregulated proteins were also implicated in many biological processes (Figure S6B and Table S4). 155 proteins were involved in metabolism, including 39 in lipid transport and metabolism. 44 were related to signal transduction and mechanisms. The most markedly reduced proteins included Surf4 and proteins involved in lipid metabolism, such as Cyp2b10, Cyp3a25, Ephx2, Cyp2d26, Ennp7, Cyp2d26, Rdh7, Cyp27a1, apoA4, and apoC3. We also observed a significant reduction in proteins involved in immune response (Ifit1, Isg15, Enp3), glucose uptake (SLC2A2), and cholesterol absorption (NPC1L1) (Table S4). Taken together, these findings demonstrate that Surf4 deficiency causes changes in diverse biological processes in mouse jejunum.

### Effect of intestinal Surf4 knockdown on serum lipoprotein metabolism

Our next experiments were to investigate the effect of intestinal Surf4 silencing on serum lipid levels. Mice were fasted and then received an oral gavage of corn oil. Male but not female *Surf4*^IKO^ mice displayed a downward trend in fasting serum TG levels than corresponding flox mice, but the difference did not reach statistical significance (Figures 6A and B). On the other hand, fasting serum FFA (6C and D), TC (6E and F) and HDL-C (6G and H) levels were all significantly reduced in both male and female *Surf4*^IKO^ mice. After a lipid bolus, serum TG, TC, FFA and HDL-C levels were all dramatically increased in the re-fed *Surf4*^flox^ mice compared to the fasted *Surf4*^flox^ mice, whereas post-fat feeding serum TG, TC, and HDL-C levels were not significantly increased in the re-fed *Surf4*^IKO^ mice compared to the fasted *Surf4*^IKO^ mice. *Surf4*^IKO^ mice did show a significant increase in serum FFA levels after fat-feeding, but to a lesser extent than *Surf4*^flox^ mice (Figures 6C and D). These findings were consistent with data in the [^14^C]-oleic acid experiment (Figure 4G) and further indicate insufficient dietary lipid absorption and secretion in *Surf4*^IKO^ mice. We also measured non-fasting serum lipid levels. Compared to *Surf4*^flox^ mice, male and female *Surf4*^HTZ^ mice had comparable serum TG, TC, and HDL-C levels, whereas male and female *Surf4*^IKO^ mice showed a significant reduction in serum TG, TC, and HDL-C levels, although the decrease in serum TG levels in female *Surf4*^IKO^ mice was smaller than that in male *Surf4*^IKO^ mice (Figures S7A to S7C). In addition, *Vil*1Cre-*Surf4*^flox^ mice injected with corn oil exhibited comparable serum TG and TC levels as *Surf4*^flox^ mice (Figures S7D and S7E). Fractionation of serum lipoproteins by FPLC also revealed a reduction in TG and TC in male and female *Surf4*^IKO^ mice (Figures S7F to S7I). Therefore, Surf4 deficiency in the intestinal epithelium reduces serum lipid levels, especially in the re-fed state.

**Figure 6.**
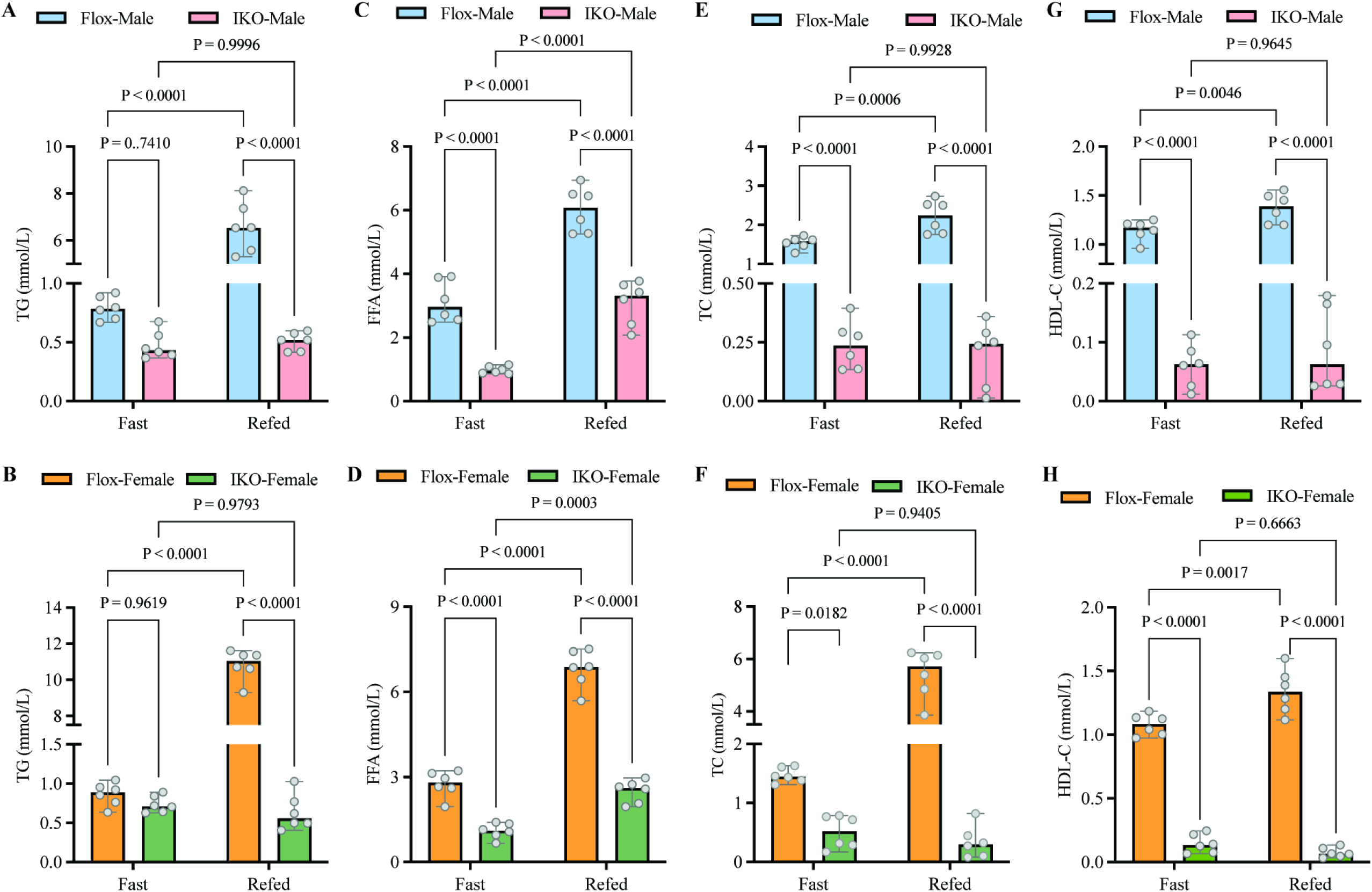
Effect of intestinal Surf4 deficiency on serum lipid levels. **A-H.** Fasting (Fast) and post-fat feeding (Refed) serum lipid levels. Male (top panel) and female (bottom panel) *Surf4*^flox^ and *Surf4*^IKO^ mice (9-10 weeks old, n=6, biological replicates) were injected with tamoxifen for 4 days, and fed a standard laboratory rodent diet for 4 days. Mice were then fasted for 10 h and received an oral gavage of olive oil. Blood samples were collected before and 4 h after oral gavage. Fasting and post-fat feeding serum levels of TG (A and B), FFA (C and D), TC (E and F), and HDL-C (G and H) were measured. Two-way ANOVA followed by a post-hoc test was used to determine the significance among different groups.

We then assessed serum lipoprotein levels and found that the levels of apoB-100, apoB-48, apoE, apoA-I, and apo-IV were all reduced in both fasted and fat-refed male and female *Surf4*^IKO^ mice (Figures 7A, 7B, and S8), indicating altered lipoprotein metabolism in *Surf4*^IKO^ mice. To further confirm these findings, we performed label-free proteomics analysis of serum samples from male mice. A total of 5551 serum peptides were identified, matching 5176 unique proteins. Of these, 912 quantifiable proteins were identified, including 170 up-regulated and 156 down-regulated proteins (Figures 7C and 7D, Tables S5 and S6). The most notable downregulated proteins in *Surf4*^IKO^ mice were apolipoproteins, such as apoA1, apoE, apoC1 and apoC2, that can be secreted from the intestine. Serum apoB levels were also significantly reduced in *Surf4*^IKO^ mice (IKO/Flox=0.746, p=3.76344E-08) (Table S5).

**Figure 7.**
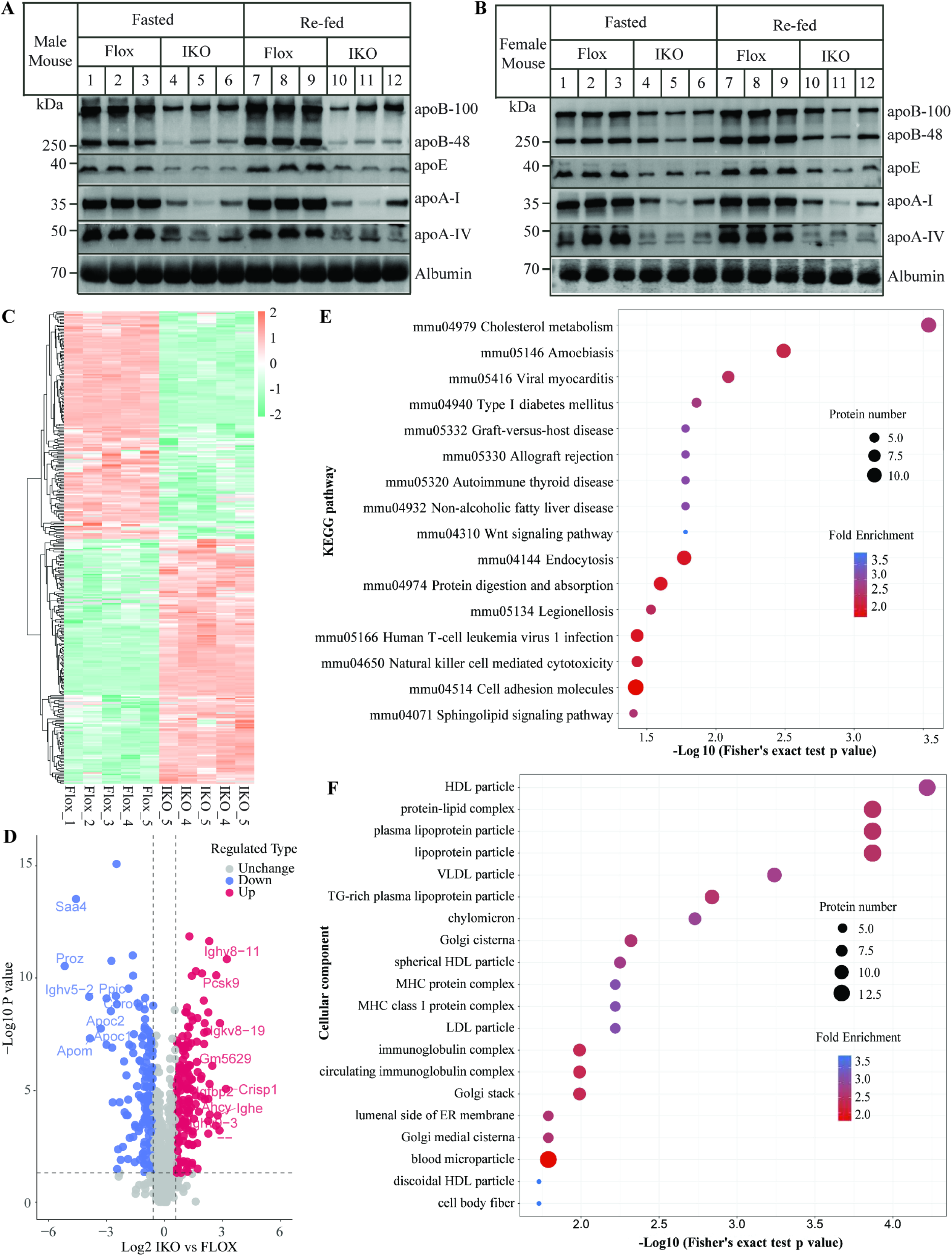
Effect of intestinal Surf4 deficiency on serum protein levels. **A and B.** Immunoblotting. Male and female mice (n=3, biological replicates) were treated as described in Figure 6 legend. Same amount of fasting (Fast) and post-fat feeding (Re-fed) serum samples was applied to immunoblotting. **C-F.** Proteomics of mouse serum (n=5, biological replicates). *Surf4*^flox^ and *Surf4*^IKO^ mice (9-week-old, male) were injected with tamoxifen for 4 days, fed a standard laboratory rodent diet for 4 days. Serum was collected for proteomics. Heat map (C), volcano plot (D), and KEGG pathway enrichment (E and F). In heatmaps, each block represents a certain protein, and the horizontal axis represents each sample. Heatmap presents differentially expressed proteins. Values were z-score normalized, log2 label-free quantification intensity. The color gradient is from green (low values) to orange (high values) Proteomic data were analyzed, and graphic were generated by online system of PTM-Bio Shiny Tool.

On the other hand, the levels of several immunoglobulin heavy chain proteins were elevated in *Surf4*^IKO^ mice (Table S6), suggesting activated immune response. We also observed significantly elevated expression of proprotein convertase subtilisin/kexin type 9 (PCSK9, IKO/Flox=6.352, p=7.694E-11) and adenosylhomocysteinase (AHCY, IKO/Flox=5.038, p=0.00012). PCSK9 is upregulated by SREBP2 at the transcriptional levels ^54^, and circulating PCSK9 is primarily secreted from hepatocytes ^55^. These indicate that the transcriptional activity of SREBP2 may be increased in the liver of *Surf4*^IKO^ mice. AHCY is ubiquitously expressed and hydrolyzes S-adenosylhomocysteine to adenosine and L-homocysteine. It is primarily localized to the cytosol and has been implicated in regulating cellular homeostasis and tissue damage ^56^. How intestinal Surf4 deficiency led to elevated serum levels of AHCY and its possible pathophysiological role are unclear. Nevertheless, KEGG pathway analysis revealed that proteins with significantly altered expression were enriched in the cholesterol metabolism pathway, followed by amoebiasis in *Surf4*^IKO^ mice (Figure 7E). Amoebiasis is an intestinal illness caused by the protozoan parasite *Entamoeba histolytica* ^57^, suggesting that Surf4 deficiency may cause intestinal damage. Compartment complex enrichment analysis showed that proteins with significantly altered expression were enriched on lipoprotein particles and protein-lipid particles, such as HDL, VLDL, and CMs, in *Surf4* ^IKO^ mice (Figure 7F). Taken together, these findings indicate that silencing intestinal Surf4 significantly reduces serum lipoprotein levels, suggesting its important role in regulating lipoprotein metabolism.

### Impact of intestinal Surf4 deficiency on the liver

We also investigated whether changes in serum lipoprotein and lipids levels affected lipid metabolism in the liver of *Surf4*^IKO^ mice. As shown in Figures 8A and 8B, the liver size and weight of *Surf4*^IKO^ mice were significantly reduced. Intestinal Surf4 deficiency also significantly reduced hepatic TG but not total cholesterol levels in non-fasted mice (Figures 8C and 8D). Serum AST activity was comparable in the two genotypes, but ALT activity was significantly increased in *Surf4*^IKO^ mice, indicating mild liver damage (Figures 8E and 8F). Analysis of mRNA levels revealed a significant increase in mRNA levels of *Srebf1c* and its target gene *Fasn*, as well as *Srebf2* and its target gene *Hmgcr* (Figure 8G), suggesting increased *de novo* lipogenesis and cholesterol biosynthesis. We then challenged mice with fasting, followed by an oral gavage of glucose. Fasting serum ketone body levels and liver TG and TC levels were lower in *Surf4*^IKO^ mice than in *Surf4*^flox^ mice (Figures 8H to 8J). 4 h after glucose refeeding, serum ketone body levels and hepatic cholesterol levels were not significantly altered, but hepatic TG levels were significantly reduced in *Surf4*^flox^ mice. In contrast, serum ketone body levels and liver TG and TC levels were increased in re-fed *Surf4*^IKO^ mice compared with fasted *Surf4*^IKO^ mice, reaching levels essentially comparable to that in re-fed *Surf4*^flox^ mice (Figures 8H to 8J). This indicates that glucose feeding could restore hepatic lipid levels in *Surf4*^IKO^ mice. We also measured serum TG and TC and found that glucose did not significantly affect serum TG levels but partially restored serum TC and HDL-C levels in *Surf4*^IKO^ mice (Figure S9). The liver may secrete some biosynthetic cholesterol into circulation to compensate for the loss of dietary cholesterol.

**Figure 8.**
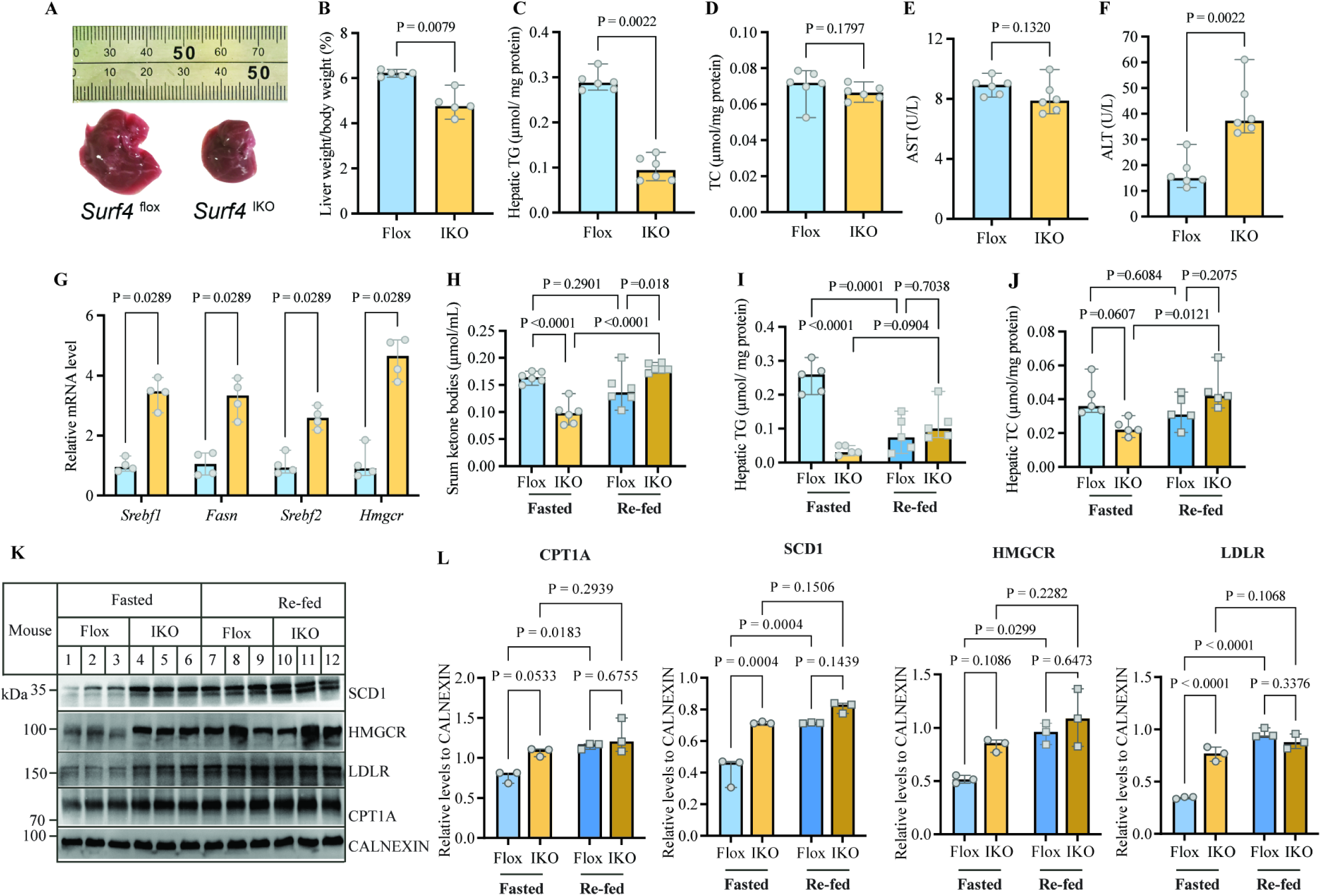
Effect of intestinal Surf4 deficiency on hepatic lipid metabolism. Nine-week-old male mice were injected with tamoxifen for 4 days and then fed a standard laboratory rodent diet for 4 days. **A.** Pictures of the liver of non-fasted *Surf4*^flox^ and *Surf4*^IKO^ mice. **B.** Percentage of liver weight to body weight (n=5, biological replicates). **C and D.** Liver lipid levels (n=6, biological replicates). Liver was collected from non-fasted male *Surf4*^flox^ and *Surf4*^IKO^ mice. Total lipids were extracted for measurement of TG and TC. **E and F.** AST and ALT of non-fasting serum (n=6, biological replicates). **G.** qRT-PCR. Total RNA was extracted from the liver of non-fasted mice (n=4, biological replicates). **H-L.** Glucose refeeding. Male *Surf4*^flox^ and *Surf4*^IKO^ mice were fasted for 10 h. Half of mice received an oral gavage of glucose solution (1g/kg, refed group). 4 h later, fasted and refed mice were euthanized. Serum ketone (n=6, biological replicates, H) was measured. Total lipids were extracted from the liver (n=5, biological replicates) and subjected to measurement of TG (I) and TC (J). Equal amount of liver homogenate was subjected to immunoblotting using antibodies indicated (K). Densitometry was determined. Relative expression was calculated as the ratio of the densitometry of targets to that of CALNEXIN (L). The Mann-Whitney test was used to determine significant differences between two groups of panels B-G. Two-way ANOVA followed by a post-hoc test was used for statistical analysis of panels H-J and L.

We then assessed the expression of several proteins involved in lipid metabolism. SCD1 and HMGCR are rate-limiting enzymes in *de novo* lipogenesis and cholesterol biosynthesis, respectively. Hepatic LDLR is primarily responsible for clearance of circulating LDL and CM remnants ^58, 59^. CPT1A plays a critical role in fatty acid β-oxidation. Fasting and refeeding did not significantly alter CPT1A levels in both *Surf4*^flox^ and *Surf4*^IKO^ mice, although they all showed an increasing trend in the re-fed mice. On the other hand, glucose refeeding significantly increased the levels of CPT1A, SCD1, HMGCR and LDLR in the liver of *Surf4*^flox^ mice (Figures 8K and 8L), suggesting increased fatty acid β-oxidation, *de novo* lipogenesis and cholesterol biosynthesis, which is consistent with previous reports^60, 61^. Compared to fasted *Surf4*^flox^, fasted *Surf4*^IKO^ mice showed a significant increase in SCD1 and LDLR, as well as an increasing trend in CPT1A and HMGCR levels. Glucose refeeding did not significantly increase the levels of these proteins in *Surf4*^IKO^ mice compared with fasted *Surf4*^IKO^ mice, although they showed an increasing trend, likely due to their elevated expression in the fasting state. Taken together, these findings suggest that *Surf4*^IKO^ mice appear to have increased hepatic lipogenesis to compensate for the loss of dietary lipids.

## Discussion

Hepatic Surf4 plays an important role in lipoprotein metabolism via facilitating VLDL secretion ^23, 35, 36^. Surf4 is also expressed in enterocytes. Here, we found that lipid absorption and secretion were significantly impaired in both male and female *Surf4*^IKO^ mice, indicating the important role of intestinal Surf4 in lipid absorption and secretion. On the other hand, we observed extensive subepithelial spaces in the jejunum of *Surf4* ^IKO^ mice even though Surf4 deficiency did not cause notable leaky intestine, indicating intestine damage. Meanwhile, we found marked and significant accumulation of TG in the intestine of *Surf4*^IKO^ mice compared to *Surf4*^flox^ mice despite impaired lipid absorption. Furthermore, unlike wild-type enterocytes that contained many CM-like vesicles, Surf4-deficient enterocytes lacked these CM-like vesicles and instead contained many small lipid vacuoles in the ER lumen. Surf4 also colocalized with apoB and co-immunoprecipitated with apoB 48 in Caco-2 cells. Therefore, our findings indicate that Surf4 facilitates CM secretion. It is most likely that the retention of small lipid vacuoles in the ER lumen of Surf4-deficient intestine results in ER stress and subsequent unfolded protein response, which causes enterocyte damage and further impairs lipid absorption and secretion. Indeed, proteomics data revealed that protein turnover and chaperons were one of the pathways enriched with upregulated proteins in the jejunum of *Surf4*^IKO^ mice. In addition, proteomics data also showed altered expression of proteins involved in diverse biological processes in *Surf4*^IKO^ mice. The most well-characterized function of Surf4 is to act as a cargo receptor to facilitate ER export of secretory proteins via a vesicle or tubular transport system. However, none of the up- and down-regulated proteins in the intestine of *Surf4*^IKO^ mice are known Surf4 substrates, with the exception of apoA-I that was increased about 4-fold in *Surf4*^IKO^ mice (Supplementary table 3). It will be of interest to investigate if Surf4 directly regulates the expression of these proteins in the intestine. Nevertheless, we cannot rule out the possibility that Surf4 deficiency somehow directly causes general damage in intestinal function, which then impairs lipid absorption and secretion and consequently causes lipid accumulation and ER lumen retention of small lipid vacuoles in enterocytes.

Unlike Surf4 hepatocyte-specific knockout (*Surf4*^LKO^) mice, which were indistinguishable from *Surf4*^flox^ mice, *Surf4*^IKO^ mice had significantly reduced body weight and survival. In mice that reached the humane endpoint, white adipose tissues were essentially gone (data not shown). It is possible that *Surf4*^IKO^ mice may die from nutrient deprivation due to insufficient lipid absorption and CM secretion in the intestine.

Tamoxifen-induced intestine-specific knockout of *Mttp* (*Mttp*^IKO^) also significantly reduced plasma TG and cholesterol levels and exhibited cytosolic accumulation of large lipid droplets and accumulation of TG but not cholesterol in enterocytes ^9^. However, enterocytes can secrete lipid-poor apoB48-containing lipoprotein particles ^62, 63^. ApoB48 secretion in *Mttp*-deficient enterocytes was detected in the lipid-poor particles. In addition, insufficient dietary lipid absorption was compensated by increased lipid biosynthesis in the liver of *Mttp*^IKO^ mice. These might explain the lack of body weight loss or increased mortality in male and female *Mttp*^IKO^ mice despite their stunted growth compared with control mice. We also observed a significant increase in the expression of lipogenic genes in the liver of *Surf4*^IKO^ mice, indicating that hepatic lipogenesis may also be increased to compensate for the loss of dietary lipids in *Surf4*^IKO^ mice. However, this mechanism may not adequately compensate for the loss of dietary lipids in *Surf4*^IKO^ mice, especially male *Surf4*^IKO^ mice. Additionally, Surf4 has been implicated in facilitating ER export of diverse secretory proteins via the canonical COPII vesicles or a tubular trafficking network. Proteomics data showed that the expression of 51 proteins with unknown functions was upregulated in the jejunum of *Surf4*^IKO^ mice. Therefore, we cannot exclude the possibility that Surf4 may mediate secretion of unidentified proteins that are critical for mouse survival.

Notably, the reduction in body weight loss and survival was significantly milder in female *Surf4*^IKO^ mice than in male *Surf4*^IKO^ mice. Mounting evidence indicates that endogenous sex hormones have different effects on lipoprotein metabolism ^64–66^. Premenopausal women have a significantly lower risk of atherosclerotic cardiovascular disease (ASCVD) than men. Studies have shown that enhanced catabolism but not reduced production of triglyceride-rich lipoprotein (TRL) particles contributes to the lower risk of ASCVD in premenopausal women ^64, 65^. Similarly, the clearance of TRL is greater in female Sprague-Dawley rats ^66^. It is of note that a vast number of studies focus on the liver and apoB100 lipoprotein metabolism due to the relatively high expression of estrogen receptors in the liver and the positive association between plasma apoB100 levels and the risk of ASCVD. Estrogen receptors are also expressed in the intestine and play important physiological roles, such as regulating water and electrolyte metabolism ^67^. However, the role of estrogen in CM secretion remains elusive. The knockdown efficiency of Surf4 in female mice appeared to be less than that in male mice. In addition, Surf4 protein levels in both male and female *Surf4*^IKO^ mice were markedly reduced compared to that in *Surf4*^flox^ mice. We found that, like male *Surf4*^IKO^ mice, female *Surf4*^IKO^ mice showed a dramatic reduction in intestinal lipid absorption and secretion, as well as post-fat feeding serum levels of TG and TC. We also observed significant lipid accumulation in the intestine of female *Surf4*^IKO^ mice. These findings suggest that factors other than impaired dietary lipid absorption and secretion may contribute to the substantial difference in body weight and mortality between male and female *Surf4*^IKO^ mice.

Women are known to survive better than men during severe famines ^68, 69^ . It has been proposed that evolution is at least partially responsible for this difference. When the food supply is short, mammalian females are under more selective pressure than males, so females are more pressured to evolve mechanisms to facilitate survival during famines. Similarly, studies in C57BL/6 mice showed that female mice survive better than male mice under prolonged fasting or starvation, probably mainly due to estrogens, which can increase fatty acid oxidation, ketone body production and heat generation to support energy metabolism and homeostasis needed for survival ^70^. We did observe a significant reduction in heat production in male but not female *Surf4*^IKO^ mice. In addition, Du et al. reported during starvation, neurons from male mice are more prone to autophagy and die, whiles neurons from females survive longer since these neurons can accumulate triglycerides and form lipid droplets ^64^. In our study, TG absorption and secretion in the intestine of both male and female mice were significantly impaired, which may result in prolonged fasting or starvation. This may explain why female *Surf4*^IKO^ mice survive better.

Mice lacking apoB in the intestine and expressing human apoB only in the liver also exhibited significantly impaired dietary lipid absorption and CM secretion ^12^. These mice were normal at birth, but developed significant growth retardation and had small liver and lipid accumulation in the small intestine. Like *Surf4*^IKO^ mice, more than half of neonatal mice lacking intestinal apoB died within 3 weeks of age, but unlike *Surf4*^IKO^ mice, they showed no sex differences. Furthermore, surviving male and female mice gained weight rapidly, eventually reaching body weights similar to their wild-type littermates, although their intestinal fat absorption remained insufficient. Plasma cholesterol but not TG levels were reduced in the adult mice lacking intestinal apoB. Apparently, these surviving mice developed a mechanism to compensate for the loss of nutrients caused by intestinal lipid malabsorption. Therefore, it is possible that male *Surf4*^IKO^ mice may not have developed a compensatory mechanism in time to make up for the loss of dietary nutrients due to knockdown of intestinal Surf4 in adulthood. Female *Surf4*^IKO^ mice, on the other hand, could survive better under starvation and eventually develop a compensatory mechanism like mice lacking intestinal apoB.

In addition, Tg(*Vil1*-cre/ER^T2^)23Syr mice were used to generate *Surf4*^IKO^ mice in our study. Compared to other *Vil1*-Cre transgenic mice, including *Vil1*^Cre/997^, *Vil1*^Cre/1000^, and Tg(*Vil1*-cre)20Syr, Tg(*Vil1*-cre/ERT2)23Syr more specifically targets the intestinal epithelium but has lesser recombination activity compared to other *Vil1*^Cre^ lines ^71^. Unlike the other lines that can drive detectable Cre expression in extraintestinal tissues, such as the liver, kidney, bladder, and stomach, the Cre activity of *Vil1*-cre/ERT^2^ is only detected in the small and large intestines of female mice. In male mice, scattered Cre activity was also detected in the testis. Consistently, we found that the mRNA level of Surf4 was mildly but significantly reduced in the testis of male *Surf4*^IKO^ mice but not in the ovary of female *Surf4*^IKO^ mice. Surf4 is a cargo receptor located in the ER membrane and facilitates ER export of secretory proteins. Its substrates have not been completely identified. Experiments are underway to investigate the function of testicular Surf4 and whether reduced testicular Surf4 expression contributes to the dramatic effects on body weight and mortality in male *Surf4*^IKO^ mice.

We also observed a mild liver injury in *Surf4*^IKO^ mice. The liver and intestine communicate with each other via the gut-liver axis ^72, 73^. We did not detect notable leaky intestines in *Surf4*^IKO^ mice. However, detailed analysis revealed a mild intestinal injury, which likely causes low-grade intestinal barrier damage and subsequently increases translocation of endotoxin and toxic metabolites. Impaired lipid absorption may also lead to gut microbial dysbiosis and alter bacterial metabolites. These changes can affect liver function via the gut-liver axis and cause systemic inflammation. Indeed, proteomics of serum samples showed elevated levels of several immunoglobulin heavy chain proteins in *Surf4*^IKO^ mice, suggesting activated immune responses.

Lack of intestinal MTP resulted in only cytosolic lipid droplet accumulation ^74^, whereas mice without intestinal apoB had ER lumen and cytosolic lipid droplet accumulation ^22^. We also observed accumulation of cytosolic lipid droplets and retention of ER lumenal small lipid vacuoles in the enterocytes of *Surf4*^IKO^ mice. It is believed that small lipid-poor apoB48 particles fuse with ER lumenal apoB-free lipid droplets to form pre-CM in the ER lumen. The process by which PCTV buds from the ER and subsequently fuses with the Golgi apparatus is well-studied and involves multiple proteins ^4, 15–22^. However, how lipid-poor apoB48 particles fuse with apoB-free lipid droplets in the ER lumen and how pre-CMs are sorted into PCTVs are unclear. Deficiency of Surf4, a cargo receptor located in the ER exit site, resulted in ER lumenal retention of small lipid vacuoles. Surf4 was colocalized with apoB and co-immunoprecipitated with apoB-48 in differentiated Caco-2 cells, suggesting an interaction between the two proteins. It is of note that Caco-2 cells express both apoB-100 and apB-48. Unfortunately, our custom-made or commercially available anti-Surf4 antibodies failed to immunoprecipitate Surf4 from tissue homogenate. In addition, while dietary fat is the main source of TG in enterocytes, emerging evidence indicates that *de novo* lipogenesis plays a role in TG production ^3, 75^. Dietary carbohydrates, such as glucose and fructose, can increase TG biosynthesis in enterocytes, which can be packaged into cytosolic lipid droplets or used for CM assembly. We did observe elevated expression of proteins important for lipogenesis in the intestine of *Surf4*^IKO^, suggesting increased *de novo* lipogenesis. Meanwhile, *Surf4*^IKO^ mice also showed increased expression of proteins important for fatty acid oxidation. High-fat diet feeding increases fatty acid oxidation in mouse enterocytes ^76^. It is unclear why Surf4-deficient enterocytes still increase lipogenesis when TG is already significantly accumulated inside cells. One possibility is that defective CM secretion in *Surf4*^IKO^ mice leads to nutrient deficiency, which may signal enterocytes to increase lipogenesis for producing more CM, exacerbating intestinal TG accumulation. This may, in turn, trigger fatty acid oxidation to alleviate lipid accumulation. However, increased fatty acid oxidation cannot eliminate the marked accumulation of TG caused by defective CM secretion. Further studies are warranted to elucidate the detailed underlying mechanism. Nonetheless, our study provides strong evidence for the essential role of Surf4 in intestinal lipid absorption and CM secretion. Intestinal Surf4 deficiency significantly impairs intestinal lipid absorption and secretion and results in the retention of small lipid vacuoles in the ER lumen of enterocytes. These findings indicate that while silencing hepatic Surf4 markedly reduces the development of atherosclerosis in different mouse models of atherosclerosis, including *Ldlr*^-/-^, *apoE*^-/-^, and PCSK9-overexpressing mice, without causing hepatic steatosis, highly cell/tissue-specific targeting is required for the potential therapeutic use of Surf4 inhibition.

## Acknowledgement

The authors thanked the Cellular Imaging Centre of the Faculty of Medicine and Dentistry at the University of Alberta for confocal microscopy.

## Supplemental Materials

Expanded Materials & Methods Table S1 and S2

Figures S1-S9

Major Resources Table

Data Set in Excel file format

